# A call for improving the Key Biodiversity Areas framework

**DOI:** 10.1101/2022.06.21.496934

**Authors:** Harith Farooq, Alexandre Antonelli, Søren Faurby

## Abstract

Eight percent of all land surface has been designated as “Key Biodiversity Areas” (KBAs). Since these areas were established based on only two percent of all terrestrial species estimated to exist, we ask what would happen if we used all species on Earth to identify additional KBAs. We explore this question at a global scale by using data from 64,110 species of animals and plants to identify how many areas could qualify as KBAs under current criteria. We find that between 26% and 68% of the world’s terrestrial areas can be classified as KBAs, depending on the spatial resolution. The total area from potential KBAs increases drastically as more species are assessed, suggesting that if all species were included, virtually all land surface could eventually meet the biological requirements for becoming a KBA. In addition, KBAs lack a data-driven ranking system, thus assuming that all KBAs are equally worthy protection. KBAs are intended to be areas which are both of biological importance and manageable but the current approach largely sidesteps the biological component. We make an urgent call for stricter criteria in the KBA methodology or alternative methodologies that allow for biologically robust area prioritization, help secure evidence-based investments, and support progress towards the targets under the new Global Biodiversity Framework.

## 1. Introduction

The upcoming Conference of the Parties (COP15) under the Convention on Biological Diversity is set to agree new targets under the Post-2020 Global Biodiversity Framework. Although details of those targets are at the moment of writing still under negotiation, a core element will likely be the protection of 30% of the Earth’s land and sea area by 2030. Which areas should be prioritized for protection is therefore a critical and timely matter in conservation (Andam et al., 2008; Geldmann et al., 2013; Silvestro et al., 2022). Without clear biological reasons to select areas, we risk that areas area solely selected because they are the cheapest to protect rather than because they are the most important for conservation.

As global biodiversity continues to decline (Andermann et al., 2020; Humphreys et al., 2019; Nic Lughadha et al., 2020) it is crucial to understand the strengths and weaknesses of existing methodologies for highlighting important areas for biodiversity. An important aspect of spatial prioritization is size. Areas that are too small may be insufficient to sustain viable populations (Baeza & Estades, 2010), while areas that are too big may be unmanageable. This can be due to higher probabilities of overlap with human settlements, or with different national or regional administrations. While most protected areas are of small size, the very few large protected areas (Rodrigues, Andelman, et al., 2004) are the ones that most contribute to the global coverage of protected areas worldwide (Cantú-Salazar & Gaston, 2010). Despite this disproportionate contribution, large protected areas only capture less than 3% of rare species of amphibians, birds and mammals (Cantú-Salazar & Gaston, 2010).

While societies failed to meet the Convention on Biological Diversity target of protecting 17% of global terrestrial land and inland water areas by 2020, this percentage would still fall short to effectively cover all species (Noss et al., 2012). Some estimates suggest that we may need to protect half of the globe to protect 80% of its species (Wilson, 2016). However, the pressure to increasingly delineate protected area to meet these targets could lead countries to protect the cheapest – rather than the most biologically valuable – land, including neglecting important species and ecosystems (Venter et al., 2014). Therefore, initiatives that have suboptimal thresholds in their intrinsic criteria to highlight important areas for conservation should be used with caution or revised, as they could inadvertently be used to justify the protection of less relevant areas.

Initiatives and schemes for area prioritization include the Important Bird Areas (Bird Life International, 2014), the Alliance for Zero Extinction (AZE) sites (Ricketts et al., 2005), B-Ranked sites (TNC, 2001), Important Fungus Areas (Evans et al., 2001), Important Plant Areas (Plantlife International, 2004), and Prime Butterfly Areas (van Swaay & Warren, 2006). These initiatives led to the creation of one of the most widely used tools in prioritization, the Global Standards for the Identification of Key Biodiversity Areas (KBAs), hereafter the “KBA Standard”. Its aims are to identify sites that contribute significantly to the global persistence of biodiversity (IUCN, 2016).

The KBA concept was first introduced in 2004 (Eken et al., 2004) and co-developed by the World Commission on Protected Areas and the Species Survival Commission (IUCN, 2016; KBA Standards and Appeals Committee, 2020). KBAs are today the standard framework for area conservation, with increasing investments in many regions around the world to support their formal protection (eg. Bezos Earth Fund: https://www.birdlife.org/news/2021/12/06/bezos-earth-fund-bets-on-birds/). The KBA Standard uses a system of criteria for identifying whether a site qualifies as a KBA, based on criteria such as presence and proportional inclusion of threatened species and ecosystems, species’ distribution ranges, ecological integrity and irreplaceability. Unlike most of the aforementioned programs, KBAs have been extended to apply to any species (Eken et al., 2004; IUCN, 2016; KBA Standards and Appeals Committee, 2020).

Since KBAs build on methodologies that were constrained either by taxonomic group (e.g. Important Bird Areas, Important Plant Areas), restricted distributions (e.g Alliance for Zero Extinction), or only applied in some regions (e.g. B-Ranked sites), there are two potential pitfalls that could have been created through this aggregation and which have not yet been thoroughly investigated. Firstly, the fact that a single species is enough to trigger the KBA status of a site, and secondly, that there is no upper limit to the size of KBAs (KBA Standards and Appeals Committee, 2020).

The first problem is of particular concern, and the one we focus on here. Out of the 6.5 million terrestrial species that are expected to exist (Mora et al., 2011), (with other estimates reaching up to 6 billion (Larsen et al., 2017)) we have only described approximately 2.1 million species and assessed the conservation status of around 160,000 species (IUCN, 2020) – a minute percentage of all species to occur on the planet. This number, although small, has been used as a basis for the identification of over 11,000 KBAs (BirdLife International, 2020b), covering over 8% of the planet’s land. With some 18,000 species being described as new to science each year (IISE, 2011), some of those will inevitably provide the evidence for the creation of new KBAs. This urges the question: will there be any non-KBA area left on the planet once we assess the conservation status of all species?

Beyond biodiversity data, the KBA methodology aims at being a bottom-up approach that also takes into consideration other aspects not readily integrated in analytical frameworks. After a candidate KBA has passed all the biological criteria, it needs also to be considered manageable. Manageability is a concept that is agreed between the KBA assessors based on features such as accessibility, geographical features, socioeconomic or cultural values. This means that despite certain merits, deciding whether or not a KBA is ‘manageable’ may not constitute an objective, data-driven or reproducible decision.

Here we hypothesize that, as more species are considered when delineating KBAs, more territory meets the KBA biological requirements – a process that could continue to an extent where the biological features are no longer be relevant, and manageability become the only factor deciding whether an area should be a KBA. We focus on potential KBAs – i.e., grid cells that can in theory trigger KBA status for criteria A1a), b), e) or B1 (the biological criteria).

## 2. Methods

### Datasets

We first calculated the size distribution of existing KBAs (BirdLife International, 2020b) to design the experimental set-up of our grid-cell analyses. Since KBAs may include both terrestrial and marine areas, and we only focus on terrestrial areas here, we first identified terrestrial KBAs which we defined as as those having at least 90% of their range on land.

We downloaded global distribution ranges of 64,677 terrestrial and freshwater species from IUCN (IUCN, 2020), BirdLife International (2020a), and Roll et al. (2017), including all IUCN Red List assessed terrestrial and freshwater animals and plants. We added data from Roll et al. (2017) who presented additional maps for 3517 species of reptiles that had not yet been assessed by IUCN. We treated those species as Least Concern, as a conservative approach (i.e., to avoid identifying more KBAs than should be justifiable by available data). We expect some of these species to have a higher threat status than what we considered here, but this would only increase the number of cells identified as KBAs.

Using these range maps, we clipped the species range polygons to comprise only the areas included in terrestrial ecoregions in Olson et al. (2001). We did this to avoid triggering KBA status in sea cells, which otherwise could occur due to e.g. species with both marine and non-marine life stages, such as anadromous fish and seabirds. We removed all cells belonging to rock and ice biomes. These areas comprise glaciers and bare rock, which are generally covered by very limited, if any, vegetation cover. These clipping steps removed 0.9% of the species (567 out of 64,677).

### Gridded potential KBA maps

We then produced grids of cells that fulfil the biological criteria for being designated a KBA (hereafter, potential KBA cells) using the R package WEGE (Farooq et al. 2020). We followed the approach described in Farooq et al. (2020), where species ranges and threat status are used against the sub-criteria within two of the five main KBA criteria (A1a, A1b, A1e and B1(IUCN, 2016; KBA Standards and Appeals Committee, 2020)) to assess whether it triggers KBA status of a grid cell. A1a are sites that have ≥0.5% of the global population size and ≥5 reproductive units of a Critically Endangered (CR) or Endangered (EN) species, A1b are sites that comprise ≥1% of the global population size and ≥10 reproductive units of a VU species, A1e are sites that have effectively the entire global population size of a CR or EN species, and B1 are sites that regularly hold ≥10% of the global population size and ≥10 reproductive units of a species (IUCN, 2016; KBA Standards and Appeals Committee, 2020). Since we only use a subset of the available criteria for defining KBAs, the actual number of cells which could trigger KBA status should be even higher than the numbers we estimate. We performed all analyses at global extent in resolutions equivalent to 25 × 25 km (625 km^2^), 50 × 50 km (2500 km^2^), 100 × 100 km (10,000 km^2^) grids in a Berhmann projection.

### Number of species triggering potential KBAs

To analyze the sensitivity of KBA assignments concerning the overall species numbers, we randomly sampled a different number of species (from one to all species included in this study) 1,000 times, each time identifying how many KBAs would be inferred. Since we rely on ranges that are only expected to be accurate on a scale of around 1 × 1 degree, or roughly 100 × 100 km (Di Marco et al., 2017), there is a risk of both over and underestimates of the number of potential KBA cells. Unlike the underestimation of potential KBA cells, an overestimation could have an effect on our conclusions. To reduce this issue, we carried out three sets of supplementary analyses requiring a minimum of 1 to 5 species to be inferred in a cell for it to be a KBA. We assumed false positives (i.e. species being coded as present in a cell, although they do not occur in it) to be present in the dataset. Distributions are however generally much more carefully mapped for threatened or range-restricted species and since the KBA criteria used in this analysis focused on these, the probability of over-predicting presence of species that can trigger KBA status should be small.

## 3. Results

We retrieved information for 15,880 KBAs globally, ranging in size from < 0.0015 km^2^ to over 710,000 km^2^, with a median of 133.3 km^2^ and a mean of 1,270.2 km^2^. They cover approximately 8% of the terrestrial and 3% of the total surface area of the Earth. There are 11,879 terrestrial KBAs (Fig. 1A), of which 24.1 % are larger than 625 km^2^, 7.2 % are larger than 2,500km^2^, 1.5 % are larger than 10,000 km^2^ (Fig. 1B).

**Figure 1:**
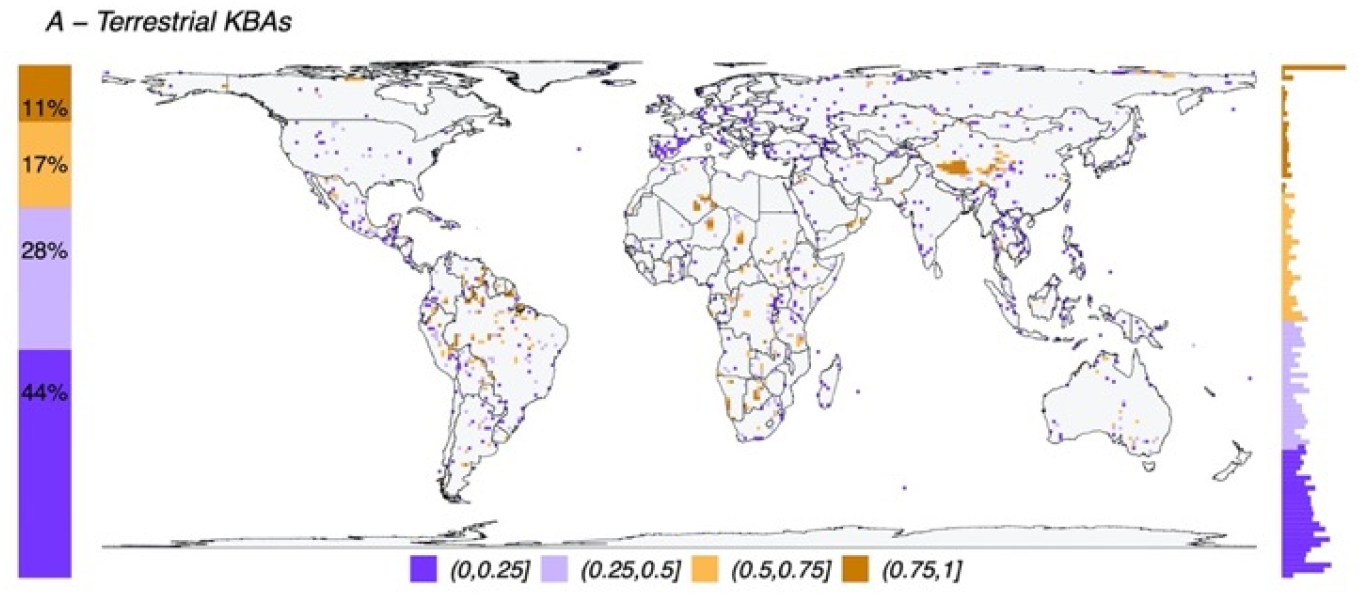
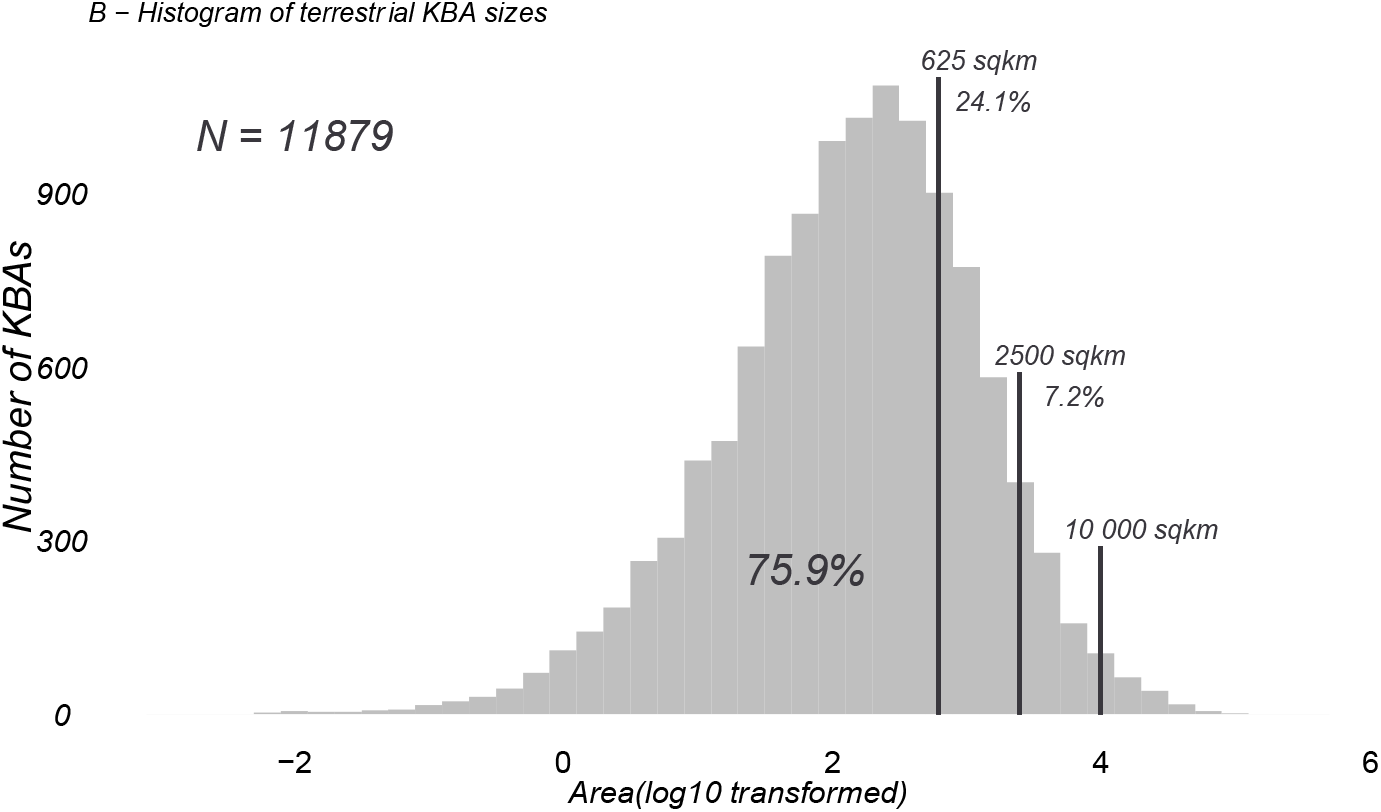
A. Global map (in a 100 × 100 km grid) showing the fraction of cells currently designated as terrestrial Key Biodiversity Areas (KBAs; N = 11,879). On the left of the map is the proportion of each grouping used in the gridded map and on the right is a histogram of the distribution of the KBA proportion from 0 in the bottom to 1 on top. B. Histogram of all existing terrestrial KBAs distributed by log10 transformed area in square kilometres. Vertical lines show where the grid sizes used in this study (625 km^2^, 2500 km^2^ and 10,000 km^2^) fit in the overall distribution of KBA sizes. Due to the relatively coarse resolution of species distributions available globally, our analyses reflect sizes above 76% of terrestrial KBAs (smallest grid size 625km^2^).

By combining all potential KBA cells obtained from our analyses, a total of between 26–68% of all terrestrial cells meet the biological requirements to be classified as KBAs (Fig. 2), depending on the resolution (26% for grids of 625km^2^, 45% for grids of 2500 km^2^, 68% for grids of 10,000 km^2^). At the highest resolution (smaller grid size) 21% of the potential KBA cells are triggered by over 5 species and 39% are triggered by just one, while at the lowest resolution (greatest grid size), 52% of the potential KBAs are triggered by over 5 species and 17% are only triggered by one species. Crucially, the more species are included in the analyses, the more potential KBAs are generated (Fig. 3).

**Figure 2:**
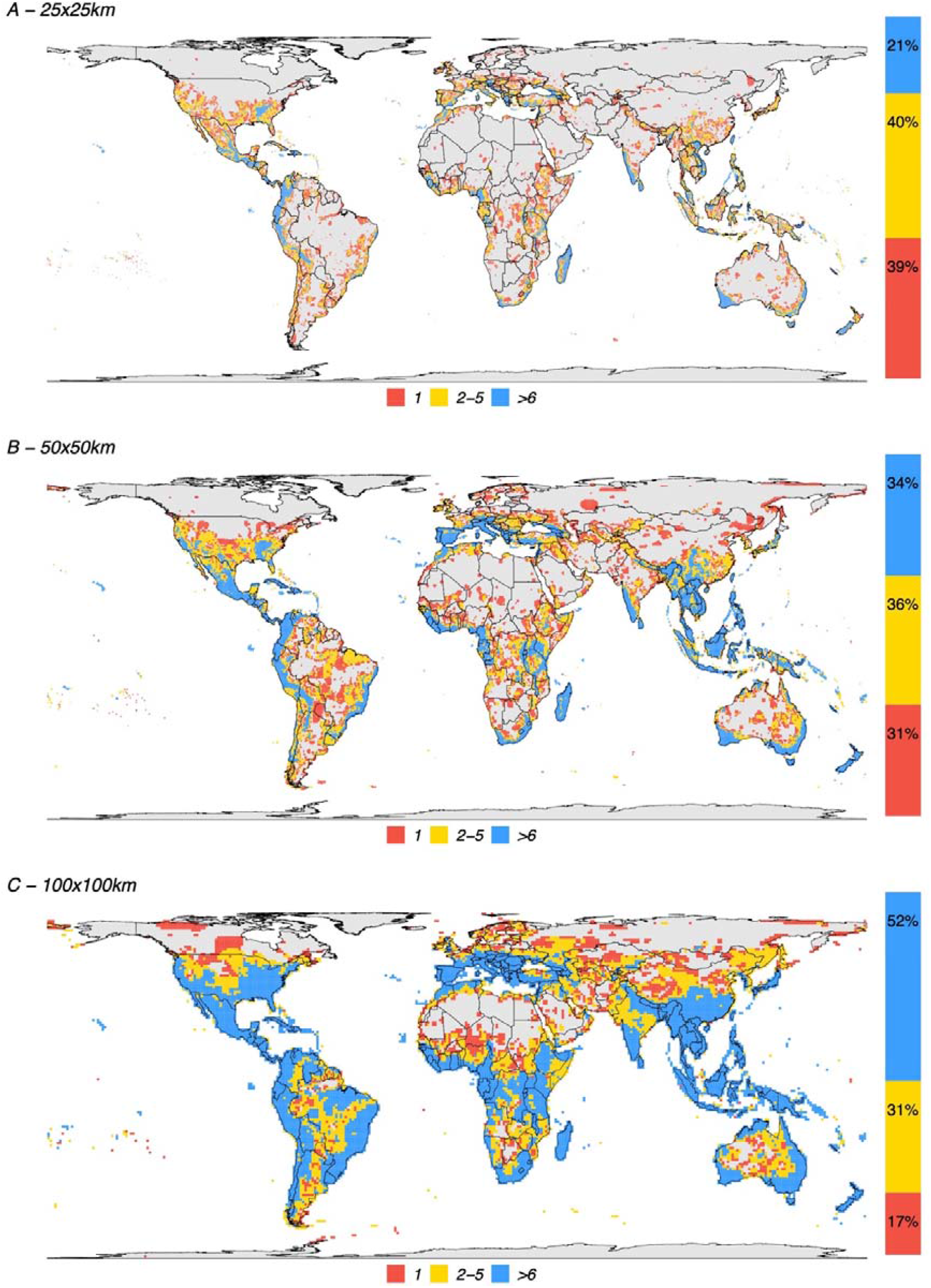
Spatial distribution of all KBA-triggering cells for all species used in this study. A. A. 25 × 25 km grid resolution, B. 50 × 50km grid resolution and C. 100 × 100km grid resolution. Fifty-five countries, including Italy, Greece, Malaysia, Haiti, Gabon, Madagascar and New Zealand, have at least 90% of their territory as potential KBAs even at the highest resolution (i.e., the smallest grid sizes of c. 25×25km).

**Figure 3.**
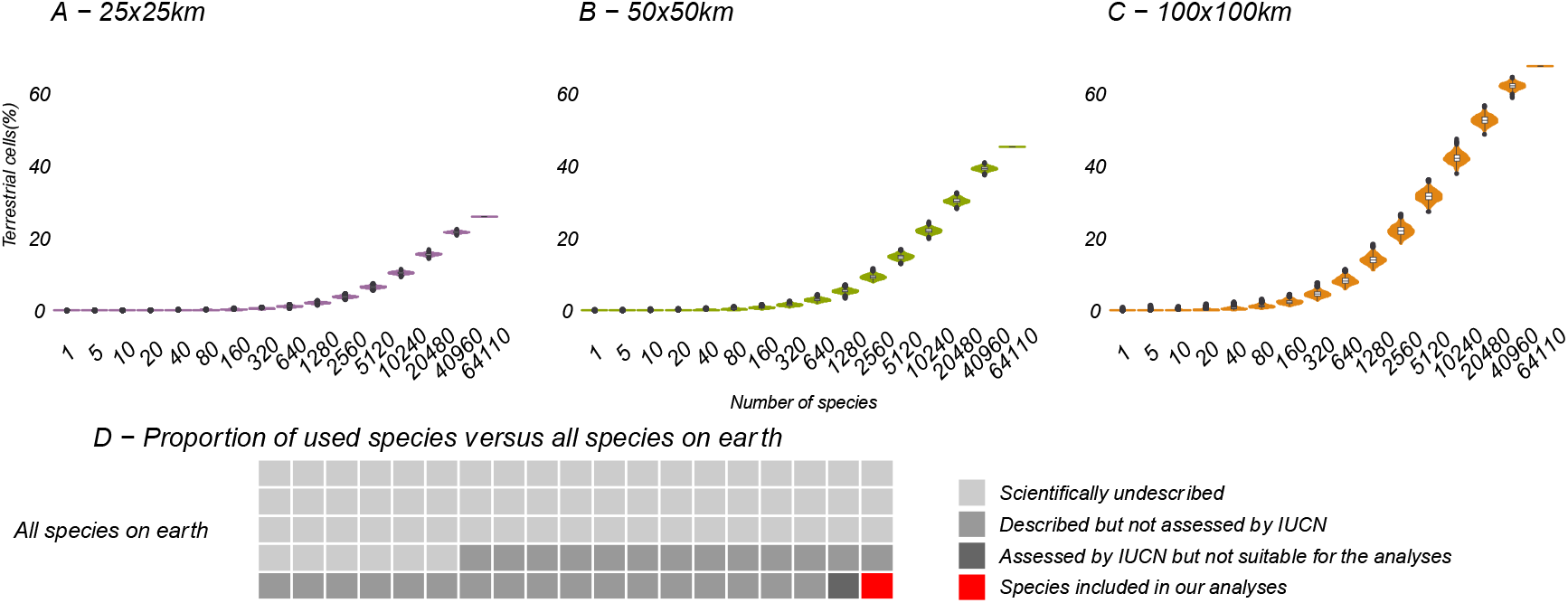
Relation between the number of species included in the analysis and potential KBA grid cells, at 25 × 25 km, 50 × 50 km and 100 × 100 km resolution: The violin plots show the number of potential KBA cells identified based on 1000 randomized subsets of each number of species. In D, the proportion of used species versus all terrestrial species expected to occur on earth, suggesting that once we use all species for the KBA assessment, virtually everywhere can become a KBA if the site is manageable. Total species: (∼6,500.000: Mora et al., 2011), described (2,115.985: IUCN, 2021), assessed (142,577: IUCN, 2021) and included in the analysis (64110).

## 4. Discussion

As we added species in the analysis, the number of potential KBAs increased steadily (Fig. 3). Given that our study was based on just 64110 species, it is plausible that if all living species had been included in this exercise, effectively any area of the world could be a potential KBA at the resolutions examined. This is a result of a combination of no upper size limit and low threshold of only one species to trigger KBA status.

Biodiversity knowledge is biased both taxonomically and spatially (Meyer et al., 2015; Troudet et al., 2017). These biases, and their associated knowledge gaps, interfere with our ability to accurately identify the most important areas to protect biodiversity. This is because we need to have information on how species are distributed and their threat status in order to map conservation priorities. We are therefore limited to accessed species by IUCN, which despite great global efforts by many voluntary experts and partner organizations, have only evaluated ca. 1% of all species estimated to exist (IUCN, 2020). The taxonomic biases in those assessments are clear: almost all described vertebrates have been assessed, contrary to the vast majority of invertebrates and fungi.

As more species are described and their conservation status is assessed, more KBAs will invariably be identified under the current classification criteria. By continually adding species to the analysis and calculating which grid cells would trigger the biological requirements for a KBA, our results shows that this process continues to the point where most terrestrial cells have at least one species to make the grid cell a potential KBA. Predictably, the coarsest resolutions show the beginnings of an asymptote due to the high proportion of required land. Importantly, the graphs at coarser resolutions (Fig. 3B,C) suggest a steep upward curve, showing that as more species are described (and assessed by IUCN), the percentage of remaining land that is not important for biodiversity will quickly diminish. The estimated number of undescribed species is over 100 times the number of species used in this study. A disproportionate number of these newly described species tend to be already under threat (Liu et al., 2022), implying that their probability of being used as basis for the identification of KBAs is also higher than the currently described and assessed species.

We have previously shown that conclusions on biodiversity patterns are highly scale-dependent (Daru et al., 2020). Here we report a similar situation concerning grid cells that trigger the biological requirements for a KBA. We find that the coarser the resolution of the analyses, the larger the fraction of cells. For the resolutions of 25 × 25 km and 50 × 50 km, 39% and 31% of the cells, respectively were triggered by the presence of just one species (Fig 2). This may likely represent a limitation of the current methodology, whereby increasing the polygon of a potential KBA, one range-restricted or threatened species will eventually overlap with the selected area.

Our conclusions do not mean that large, protected areas are not desirable. If well implemented, due to their increased connectivity, these areas will contribute to the geneflow between distant patches (Clergeau & Burel, 1997), allow for undisturbed species migrations (Ferreras, 2001) and even contribute to climate change resilience by allowing species to adjust their distributions (Opdam & Wascher, 2004). Large KBAs are often desirable for biodiversity protection and one of the criteria for KBA selection (criterion C, ecological integrity) arguably requires KBAs to be as large as the ones we focus on here. However, our results highlight that criteria A1, A1b, A1e and B1 are not realistically applicable in their current form for larger areas. By increasing the polygon of a potential KBA, a single range-restricted, or threatened species will nearly always overlap with the area and trigger a potential KBA status.

The biological component is of course only one part of the KBA selection process and the vast majority of the cells we identity could fail other aspects of KBA selection dealing with manageability. Even though we show that at 100 × 100km resolutions, a disproportionate amount of land would trigger KBA status, few if any of such areas would be manageable in practice. But crucially, if any area meets the biological requirements to become a KBA, then the KBA assessment process risks becoming solely based on land manageability, which in turn could end up neglecting the most important areas from the perspective of biodiversity outcomes and value. The greater proportion of small-sized KBAs currently recognized may demonstrate that most practitioners tend to favor smaller areas over proposing large KBAs, and therefore they are likely to already follow suggestions similar to what we propose here – but that consistency would be improved if they could be formalized.

### 4.1 Analytical limitations

By using inferred species occurrences from IUCN range maps, our analysis will most likely fall victim to commission errors – when a species is erroneously assumed to be present in an area (Rodrigues, Akçakaya, et al., 2004). However, these will be more likely associated with species with wide ranges, which in turn are unlikely to be particularly relevant for conservation purposes, because few species are both wide-ranged and threatened (Farooq et al., 2020; Keith et al., 2018).

We acknowledge that our analyses were performed on relatively large units – only 24% of current KBAs are larger than our smallest analyzed resolution of 625 km^2^ (Fig. 1). As previously mentioned, this restriction comes from the data itself, as it has been suggested that coding for species presence/absence below 1×1 degree (≈100×100 km) is often unreliable (Di Marco et al., 2017). It is therefore plausible that the trend we report may disappear at very fine resolutions and there is likely a size where the KBA Standard thresholds as currently defined would not inflate as a function of number of species included in the analysis. We cannot identify this size based on currently available data, although if it exists, it is likely substantially smaller than the smallest size we analyzed here, and would require very high-resolution species occurrence datasets not available for the majority of regions and taxa at any point soon (Farooq et al., 2021).

### 4.2 Possible solutions

One possibility to prevent the identification of so many areas as KBAs would be to impose additional restrictions for classifying an area as a KBA. This could be based, for instance, on approaches comparable to the ones used to identify biodiversity hotspots (Myers et al 2000), requiring that a KBA needs to be important for a given number of species. However, such thresholds might become arbitrary and context-dependent, making the methodology differ between regions and biomes with different amounts of biodiversity.

A better solution may be to carry out hierarchization based on continuous metrics. One option is EDGE (Evolutionarily Distinct and Globally Endangered) which ranks species according to their evolutionary distinctiveness and threat status (Isaac et al., 2007). Another is WEGE (Weighted Endemism including Global Endangerment), which weighs areas based on the conservation status and range size of the species found within them, allowing the ranking of top priorities for conservation in geographically constrained regions, such as individual countries or states (Farooq et al 2020).

Systematic conservation planning has also been proposed to prioritize between KBAs (Smith et al., 2019). This is done through the combination of biodiversity and implementation-relevant data to guide management actions based on variables such as funding, existing threat or the percentage of management targets. Additional approaches are emerging that integrate various biological and socio-economic data sources within an artificial intelligence framework, such as e.g. Conservation Area Prioritization through Artificial INtelligence (CAPTAIN: Silvestro et al., 2022).

We acknowledge that real-world decisions by policymakers will always be based on a multitude of additional aspects that go beyond biodiversity metrics, often at very fine spatial scales. Of particular relevance are the provision of ecosystem services and nature’s other contributions to people, the price of land, opportunity costs, accessibility, and conflicting interests. We are therefore by no means suggesting that decisions solely should be taken based on estimated biodiversity levels. Our key message is solely that biodiversity should be one of the major criteria and that the current KBA approach effectively removes the biodiversity component in many cases, making any large area a potential KBA.

## 5. Conclusions

This study demonstrates that the current KBA Standards are not scalable to all biodiversity and that stricter criteria, or alternative approaches, are required. Our results show that for most larger terrestrial areas, there will be at least one species capable of triggering KBA status. This is problematic because almost any site can contain a high number of micro-organisms found nowhere else (Ramirez et al., 2014; Ritter et al., 2020). If everywhere can trigger the biological requirements for a Key Biodiversity Area, then nowhere can be truly regarded as ‘key’. The fact that the effectiveness of land protection varies so much across the world, many times not meeting the intended conservation goals (Geldmann et al., 2019; Sounders et al., 1991) may be partly a consequence of the lack of hierarchization that underlies conservation. The application of formal data-driven methods should help reduce ambiguity in the selection of important areas for protection and provide a basis for better informed decisions.

## Data availability statement

The data analyzed in this study was a re-analysis of existing data, which is openly available from the references cited.

## Acknowledgements

We thank our colleague Allison Perrigo for proofreading this manuscript. S.F. and A.A. are supported by the Swedish Research Council (grants #2021-04690 and #2019-05191, respectively) and A.A. is further supported by the Royal Botanic Gardens, Kew.

